# miR-202 drives medaka fertility by targeting antagonistic Yap-dependent transcriptional regulators *tead3b* and *vgll4b* in a sex-dependent manner

**DOI:** 10.64898/2026.02.12.705488

**Authors:** Sarah Janati-Idrissi, Camille Lagarde, Aurélien Brionne, Manon Thomas, Hervé Seitz, Violette Thermes, Julien Bobe

## Abstract

How miRNAs can sometimes drive major organism-level phenotypes by targeting a single gene and by triggering limited changes in mRNA levels remains poorly understood, especially in vertebrates. In medaka, the knockout of miR-202-5p, a gonad-specific miRNA in vertebrates, leads to impaired male and female fertility, including drastically reduced egg production and low developmental success. Here we show that miR-202-5p drives gamete formation by targeting antagonist Yap-dependent transcriptional regulators *tead3b* and *vgll4b* in a sex-dependent manner. Disruption of the miR-202-5p binding site in the 3’UTR of *tead3b*, but not *vgll4b*, results in a significant decrease in female fertility. In contrast, disrupting miR-202-5p target site in the 3’UTR of *vgll4b*, but not *tead3b*, results in impaired male fertility. In females, 3D ovary imaging and RNA-seq analysis of isolated ovarian follicles revealed that disrupting miR-202-5p binding to the 3’ UTR of *tead3b* results in a polycystic ovarian syndrome (PCOS)-like phenotype and the expression of many PCOS-associated marker, including androgen signaling and estrogen metabolism genes. In males, disrupting miR-202-5p binding to the 3’UTR of *vgll4b* triggers severe phenotypes, including reduced sperm motility and abnormal testicular development. No effects on sex ratio were observed, indicating that miR-202-5p drives gamete formation by regulating mechanisms acting down-stream of the sex-determining cascade. The analysis of miR-202-5p target sites in 3’ UTRs suggests long-term conservation of antagonistic *TEAD* and *VGLL* targeting across vertebrate species, including mammals. Together, our results show that miR-202-5p drives fertility by leveraging antagonistic Yap-dependent transcriptional regulators in a sex dependent manner.

## Introduction

MicroRNAs (miRNAs) are small (approximately 22 nucleotides) non-coding RNAs that post-transcriptionally regulate gene expression through base-pairing to mRNA 3’ untranslated regions (3’ UTRs). In animals, miRNAs are involved in a variety of biological processes and striking phenotypes sometimes result from the deletion of a single miRNA gene (1, 2). Existing data in various species have subsequently shown that these phenotypes can be explained by the dysregulation of a single target mRNA (1, 3). The functional validation of such phenotypic targets (i.e. targets whose de-repression contributes to the phenotypic outcomes) is made by demonstrating that the miRNA gene knockout phenotypes are replicated by disrupting the miRNA target site in the 3’ UTR of a single gene (1, 2). In fact, while computationally predicted targets can give insights into the potential targeting repertoire of a miRNA, they do not provide information regarding the functional relevance of individual interactions (4, 5). To date, the number of functionally validated miRNA targets remains low (see (3, 4) for review) and limited to model species including *Cænorhabditis elegans* (6–9), *Drosophila melanogaster* (10), *Mus musculus* (11–16), and *Oryzias latipes* (17).

The importance of miRNAs in the regulation of major biological function is substantiated by several examples of striking organism-level phenotypes resulting from the knock-out of miRNA genes or clusters. It is noteworthy that these examples include phenotypes linked to the reproductive success, a key component of the life cycle that is critical for species survival. In worms, *let-7*, the first identified miRNA, is required for the vulval-uterine development and its knock-out results in lethality (7, 18, 19). Similarly, the *mir-35-41* cluster regulates germ cell proliferation, sex determination and fecundity (20, 21). In flies, the BX-C miRNA targets homothorax (*hth*) to regulate mating behavior, a critical element of the reproductive success (10). In mice, the deletion of a single microRNA cluster, *miR-17∼92*, in XY individuals induces complete primary male-to-female sex reversal (22) and the derepression of a single miRNA target causes female infertility (16). In medaka, a model fish species with a XX/XY sex determining system, deletion of the *mir-202* gene results in impaired reproductive success with reduced fertility in both males and females (23). In females, miR-202-5p regulates egg production (*i*.*e*. fecundity) by targeting *tead3b*, a Yap-dependent transcriptional regulator acting downstream of several signaling pathways, including Hippo, Notch, Wnt, TGF-b, and GPCR (24). However, the mechanisms by which a single miRNA targeting a single mRNA can trigger such a dramatic decrease in the number of eggs produced at each reproductive cycle, and overall, throughout life, remain unclear. It also remains unknown if the Tead3b/Yap axis is also targeted by miR-202 in males, or if different mechanisms are at stake to regulate fertility. Indeed, miR-202-5p is predominantly expressed in the gonads in both males and females in fish, amphibians and mammals (23, 25–29). This pattern is also consistently observed in holostean and teleostean fishes according to the FishmiRNA database (30), suggesting a long-term conservation of this gonad-predominant expression pattern.

Here we show that miR-202 drives gamete formation and ultimately reproductive success by targeting antagonistic Yap-dependent transcriptional regulators *tead3b* and *vgll4b* in a sex-dependent manner. While disruption of miR-202-5p binding site in *tead3b* 3’UTR results in a significant decrease in female reproductive success, including egg production, it has no impact on male reproduction. An opposite result is observed when disrupting miR-202-5p binding site in *vgll4b* UTR with a significantly reduced fertility that is observed in males but not in females. In both cases, no effects on sex ratio were observed, indicating that miR-202 drives reproductive success in a sex-dependent manner by regulating mechanisms acting down-stream of the sex-determining cascade. By comprehensively and dynamically characterizing oocyte growth using 3D imaging we showed that disruption of the miR-202-5p target site in *tead3b* UTR results in a phenotype that resembles polycystic ovarian syndrome (PCOS). We observed a decrease in the number of large size mature ovarian follicles associated with an accumulation of smaller size follicles in both juvenile fish at the onset of their reproductive period and in adult females undergoing reproduction. Using RNA-seq in isolated ovarian follicles, we showed that this major reproductive phenotype is associated with the dysregulation of key ovarian genes linked to PCOS, estrogen-metabolism, androgen signaling, and ovulation. In males, disruption of miR-202-5p binding in *vgll4b* UTR results in severely impaired male reproduction that can be explained by impaired sperm formation, lower sperm motility, and lower sperm viability. The analysis of miR-202-5p target sites in 3’ UTRs suggests long-term conservation of antagonistic *TEAD* and *VGLL* targeting across vertebrate species. Together, our results show that miR-202 drives fertility by leveraging antagonistic Yap-dependent transcriptional regulators in a sex dependent manner in a model vertebrate species.

## Results

### miR-202 drives reproductive success by targeting antagonistic Yap-dependent transcriptional regulators *tead3b* and *vgll4b* in females and males, respectively

When disrupting miR-202-5p binding in *tead3b* 3’ UTR (Figure 1A), we observed a significant decrease in female fertility (i.e. the ability of females to produce viable offspring) with an overall decrease in developing embryos in *tead3b* UTR mutant females compared to WT females (Figure 1B). In addition to the previously documented decrease in egg production (17), we could show that this reduced fertility was explained by an increased occurrence of spawns with less than 10 eggs, and a decreased occurrence of spawning and spawns with more than 20 eggs (Figure 1C). In contrast, when *tead3b* UTR mutant males were crossed with WT females, we did not observe any significant decrease in embryonic development compared to WT, indicating that disruption of miR-202-5p binding to *tead3b* UTR does not affect male fertility (i.e. the ability of males to produce viable offspring) (Figure 1D). Finally, we did not observe any significant effect on sex ratio (p-value = 0.3778).

**Figure 1:**
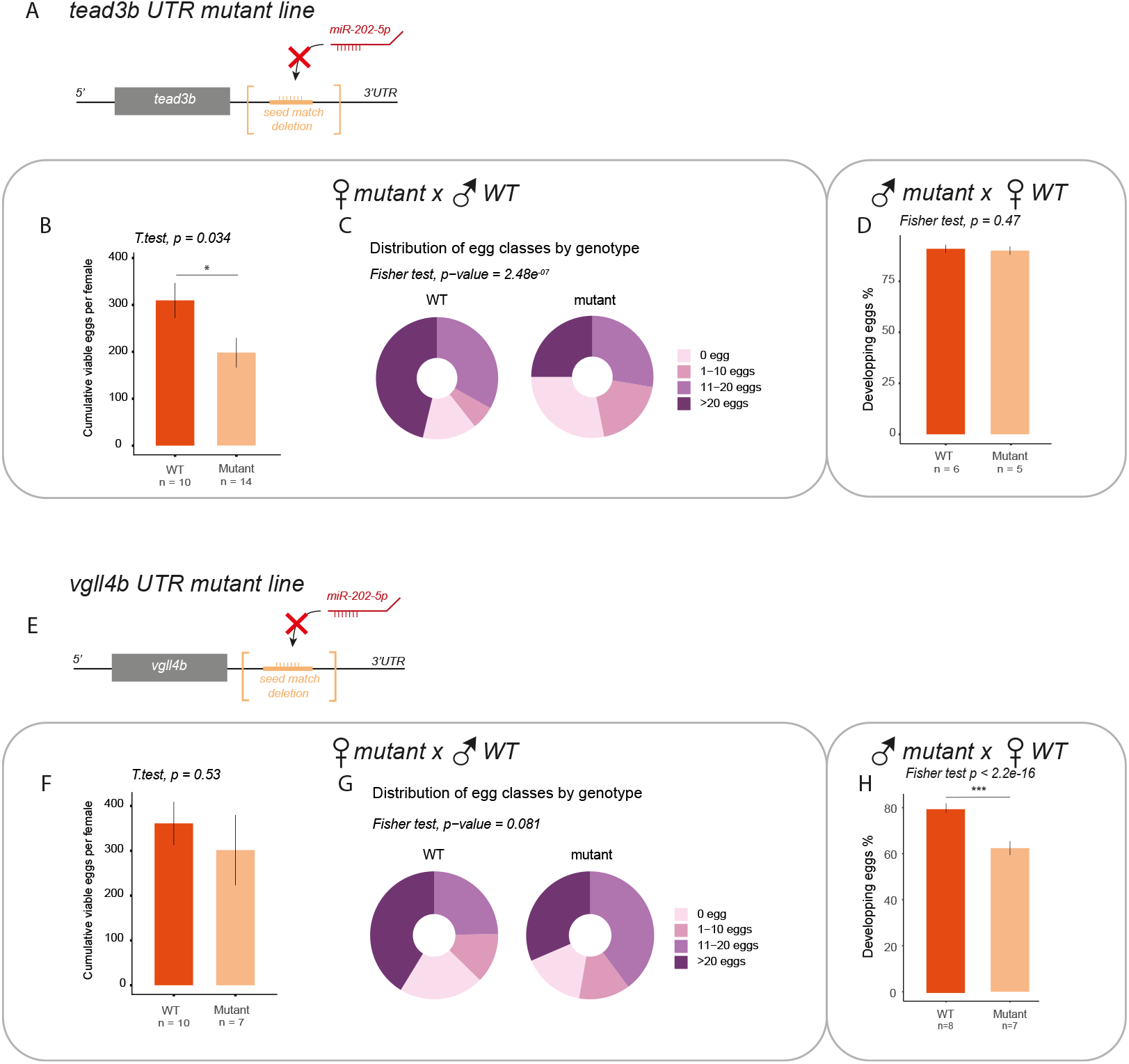
Macroscopic reproductive phenotype in *tead3b* UTR (A-D) and *vgll4b* UTR (E-H) mutant lines. For female fertility, the cumulative number of eggs leading to success full fertilization and embryonic development (B & F) and the number of eggs in each clutch (C & G) was assessed. For male fertility, the occurrence of fertilized eggs leading to successful embryonic development was monitored (D & H).

A totally opposite results was observed when disrupting miR-202-5p binding to *vgll4b* 3’UTR (Figure 1E), which had not impact on female fertility but a significant impact on male fertility. Our interest for *vgll4b* was triggered the ability of VGLL4 to inhibit YAP transcriptional activity by binding TEAD and competing with YAP in humans (31, 32) and by the presence of a 8mer miR-202-5p target site in vgll4b 3’ UTR. Mutant *vgll4b* UTR females did not exhibit any reduced developmental success when crossed with WT males (Figure 1F). Similarly mutant *vgll4b* UTR females showed spawning features, including the occurrence of large spawns, that were not significantly different from their WT controls (Figure 1G). In contrast, when mutant *vgll4b* UTR males were crossed with WT females, we observed a significant decrease in developmental success (Figure 1H). Finally, we did not observe any significant effect on sex ratio (p-value = 0.1936).

In summary, while disruption of miR-202-5p binding site in *tead3b* 3’UTR results in a significant decrease in female reproductive success, it has no impact on male reproduction. An opposite result is observed when disrupting miR-202-5p site in *vgll4b* UTR with a significantly reduced fertility that is observed in males but not in females. In addition, we did not observe any effect on sex ratio when disrupting miR-202-5p binding site in *tead3b* 3’UTR or *vgll4b* UTR, indicating that miR-202 drives reproductive success in a sex-dependent manner by regulating gamete formation through mechanisms acting down-stream of the sex-determining cascade.

### Disruption of miR-202-5p binding in *tead3b* UTR triggers a PCOS-like phenotype

To investigate further the regulation of egg formation by miR-202 we comprehensively assessed the number and size of oocytes in *tead3b* UTR mutant and WT ovaries using 3D imaging. This was performed in 3-month-old (90 days post fertilization) juveniles at the onset of their reproductive period and in 160 days post fertilization (dpf) adults during their peak reproductive period. In juveniles, we observed a dramatic decrease in the overall volume of the ovary in *tead3b* UTR mutants compared to WT (Figure 2A). When comprehensively assessing the number of oocytes in the ovary, we observed that disruption of miR-202-5p binding to the 3’ UTR of *tead3b* results in a marked accumulation of small size oocytes (i.e. less than 90 μm in diameter) and a dramatic decrease in the number of larger size oocytes (Figure 2B). This drop was especially pronounced for mid-vitellogenic (above 250 μm in diameter) oocytes that were almost completely abolished (Figure 2B-C). This shows a lower capacity of *tead3b* UTR mutant to initiate reproduction, compared to WT siblings.

**Figure 2.**
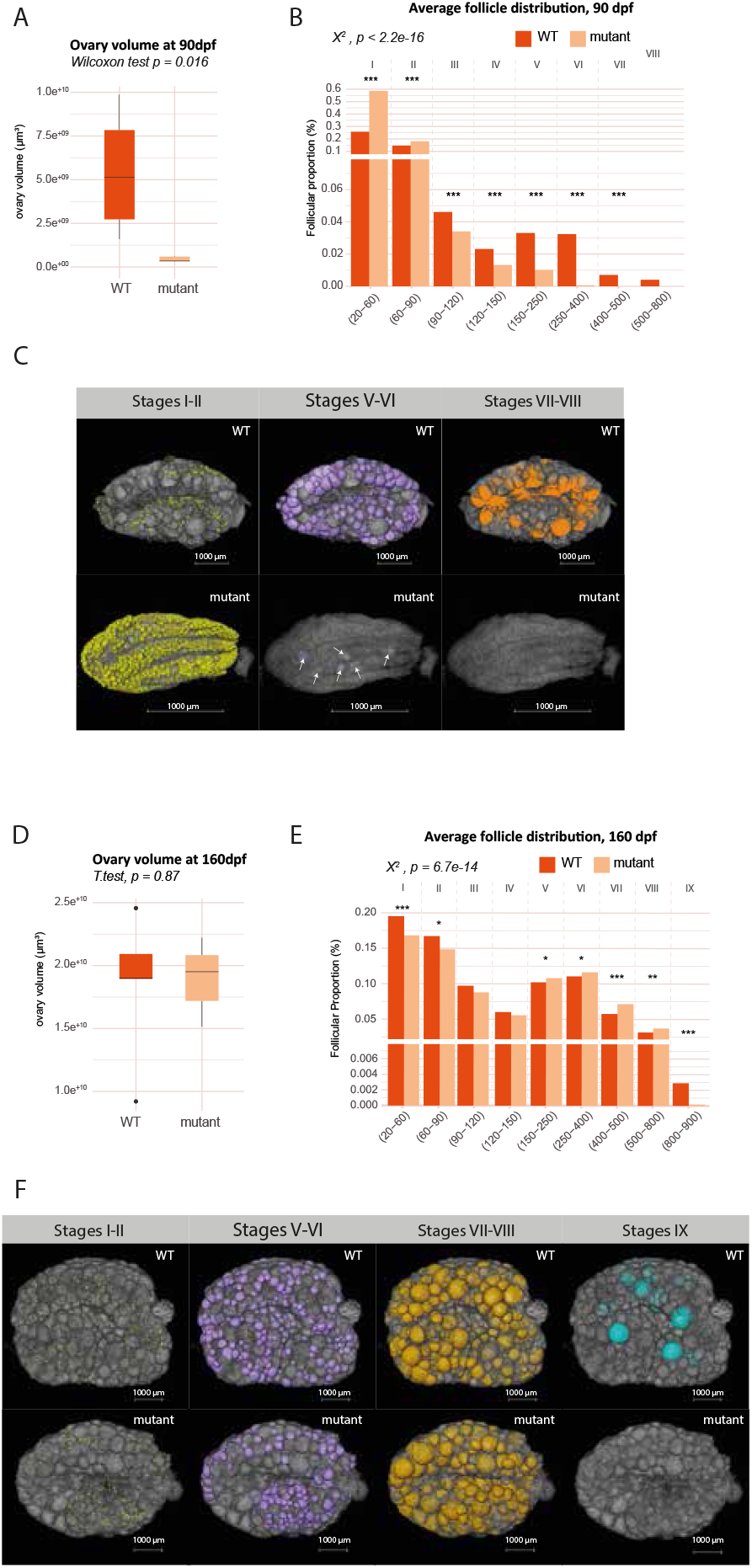
Comprehensive 3D analysis of ovarian follicle distribution in juveniles (90 dpf) (A-C) and adults (160 dpf) (D-E) in WT and *tead3b* UTR mutant females. Total ovary volume (A & D) and ovarian follicle of difference diameter (B & E) were quantified. Data represent proportions from n=4-6 ovaries per group. Statistical comparison: χ^2^ test (p < 0.001); *, **, *** indicate intervals with significant contribution to the overall difference (standardized residuals, p < 0.05, 0.01, 0.001). Representative 3D images of whole ovaries are shown (C & F).

In adults (160 dpf) that are actively reproducing, no significant decrease in total ovary volume was observed (Figure 2D). However, we observed a dramatic decrease in the number of post-vitellogenic oocytes (above 800 μm in diameter) that were almost completely abolished (Figure 2E-F). Consistent with this failure to progress from late vitellogenic to mature oocytes that can subsequently be ovulated (33) we observed an accumulation of vitellogenic oocytes (150-800 μm in diameter) (Figure 2E). In addition, we observed a decrease in the number of small oocytes (below 90 μm in diameter) that could reflect a lower ability to produce oocytes from oogonial stem cells (Figure 2E).

Together, these observations reveal that disrupting *tead3b* post-transcriptional regulation by miR-202 results in a dramatic decrease in the number of large size oocytes, including pre-ovulatory oocytes that are soon-to-be spawned, and an accumulation of smaller size oocytes. Together our observations fully explain the previously documented low female fertility phenotype and demonstrate that miR-202 regulates oocyte growth dynamics in the ovary. Disrupting miR-202 regulation results in a clear inhibition of final oocyte recruitment and growth, ultimately leading to reduced spawning and impaired egg production (i.e. low fecundity). This phenotype resembles the Polycystic ovary syndrome that is a common condition associated with a chronic lack of ovulation and accumulation of small antral ovarian follicles (oocytes surrounded by somatic follicular cells) that fail to progress to a dominant preovulatory follicle, causing arrested follicle growth (34).

### Disruption of *tead3b* targeting by miR-202-5p triggers the dysregulation of PCOS-associated genes

To disentangle the molecular mechanisms triggering the low fertility phenotype observed in *tead3b* UTR mutant females, we performed RNA-seq analysis in isolated ovarian follicles. Because *tead3b* UTR mutant females exhibit impaired oocyte development beyond late vitellogenesis, ovarian follicles were sampled from the same females at early vitellogenic (EV) and late vitellogenic (LV) stages. Gene expression trajectories between those key stages were subsequently analyzed for each individual female and contrasted between *tead3b* UTR mutants and WT. To capture key regulatory mechanisms, we isolated differentially expressed genes (DEGs) exhibiting a marked difference in expression between EV and LV stages in at least one of the experimental groups (*tead3b* UTR and WT). We then focused on the top 100 genes exhibiting the highest difference in LV/EV Log2 FC between the two experimental groups to identify genes with different transcriptional trajectories during follicular recruitment into late vitellogenesis (Supplemental data file 1). Among these 100 genes, we identified 4 genes (yellow dots) that were differentially expressed in WT but not in mutants and 14 genes (blue dots) that were differentially expressed in mutant but not in WT (Figure 3A). Remaining genes (red dots) differentially expressed in both groups but exhibiting distinct profiles (Figure 3A). Among the DEGs identified only in mutants, catechol-O-methyltransferase b (*comtb*) was the most striking, with a dramatic over expression during ovarian follicle growth in WT (log2FC = 2.32) that was almost totally abolished in *tead3b* UTR mutants (log2FC = 0.26) (Figure 3B). In mammals COMT is expressed in ovarian granulosa cells, where it converts catechol estrogens to 2-methoxyestradiol (2-ME2), a naturally occurring metabolite exhibiting anti-angiogenic activity (35) and implicated in regulating follicle homeostasis, atresia, and dominant follicle selection (36, 37). COMT is linked to PCOS at the ovarian level (38) and to other estrogen-dependent ovarian pathologies (39), including premature ovarian insufficiency (40). In fish, *comt* expression in the ovary is strongly increased during the reproductive cycle, with a peak expression during spawning, consistent with our observations in the WT group (41). In addition to *comtb*, our analyses also revealed that disruption of miR-202-5p binding site in *tead3b* UTR resulted in the dysregulation of other key genes associated with PCOS (Figure 3B), including androgen receptor (*ar*) (42), tet methylcytosine dioxygenase 2 (*tet 2*) and 3 (*tet3*) (43), as well as prolactin receptor (*prlra*) (44) and integrin subunit alpha 9 (*itga9*) (45). We also observed a dysregulation of gonadal soma derived factor (*gsdf*), a TGFB-family member, whose knock-out triggers a PCOS-like phenotype in zebrafish with reduced egg production and accumulation of immature ovarian follicles (46). Finally, cellular communication network factor 1 (*ccn1*), a known target of the YAP/TEAD axis in the mammalian ovary (47) exhibited a significantly increased expression when miR-202-5p was removed (Figure 3A-B). In humans, *CCN1* (previously known as *CYR61*) is sensitive to estrogens and associated with various pathologies (48).

**Figure 3.**
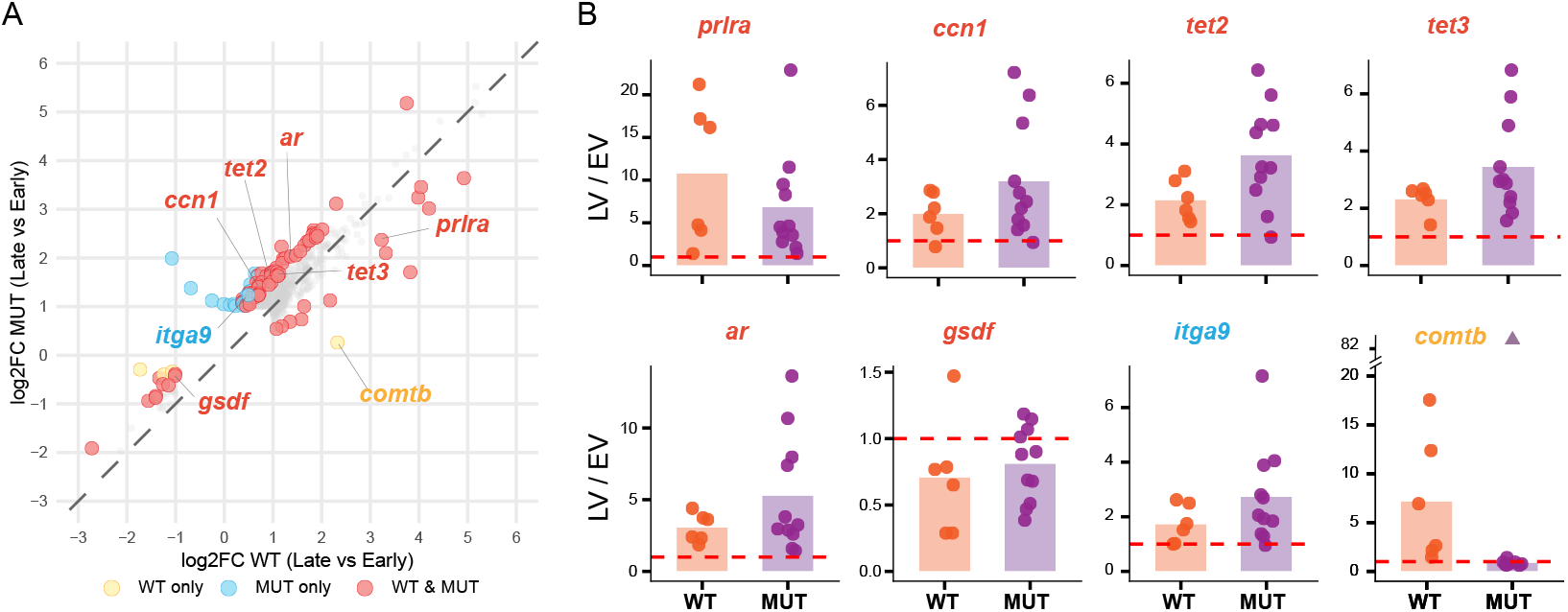
**A.** Divergent transcriptional trajectories during oocyte growth (i.e. between early vitellogenic and late vitellogenic ovarian follicles) in WT and *tead3b* UTR mutants lacking the miR-202-5p target site. **B**. Individual (dots) and mean (bars) differences in gene expression during oocyte growth in the two experimental groups. For *comtb*, outlier value in LV/EV ratio due to extremely low expression level at EV stage is displayed as a triangle and not included in mean calculation.

In summary, disruption of miR-202 target site in *tead3b* UTR leads to the dysregulation of key ovarian genes, including PCOS-associated genes, that can explain the observed impaired follicular development and overall PCOS-like phenotype. Among involved mechanisms are estrogen metabolism and epigenetic regulators that appear to play pivotal roles in PCOS.

### Disruption of miR-202 target site in vgll4b 3’ UTR triggers severely impaired male reproduction

To investigate further the reduced fertility phenotype observed in males, we characterized testicular development and sperm motility features following disruption of the miR-202 target site in *vgll4b* UTR. In vgll4b UTR mutants, we observed a highly significant decrease in both sperm motility (Figure 4A) and proportion of forward-moving sperm (progressive spermatozoa %, Figure 4B). This was associated with an increase in the proportion of slow spermatozoa (Figure 4C) and in the percentage of dead sperm (Figure 4D). Collectively, these features can explain the lower fertilization success observed when *vgll4b* UTR mutant males were crossed with WT females (Figure 1H). At the organ level, histological analyses revealed notable alterations in sperm density within mutant testes (Figure 4F). High-magnification examination of the collector canal region particularly highlighted this decreased sperm density, with mutant testes showing substantially fewer spermatozoa within the collector canals compared to the densely packed sperm observed in WT testis, suggesting impaired sperm production and/or accumulation. In summary, disruption of miR-202 target site on *vgll4b* UTR leads to severely impaired sperm production and quality that can collectively explain the reduced male fertility phenotype.

**Figure 4.**
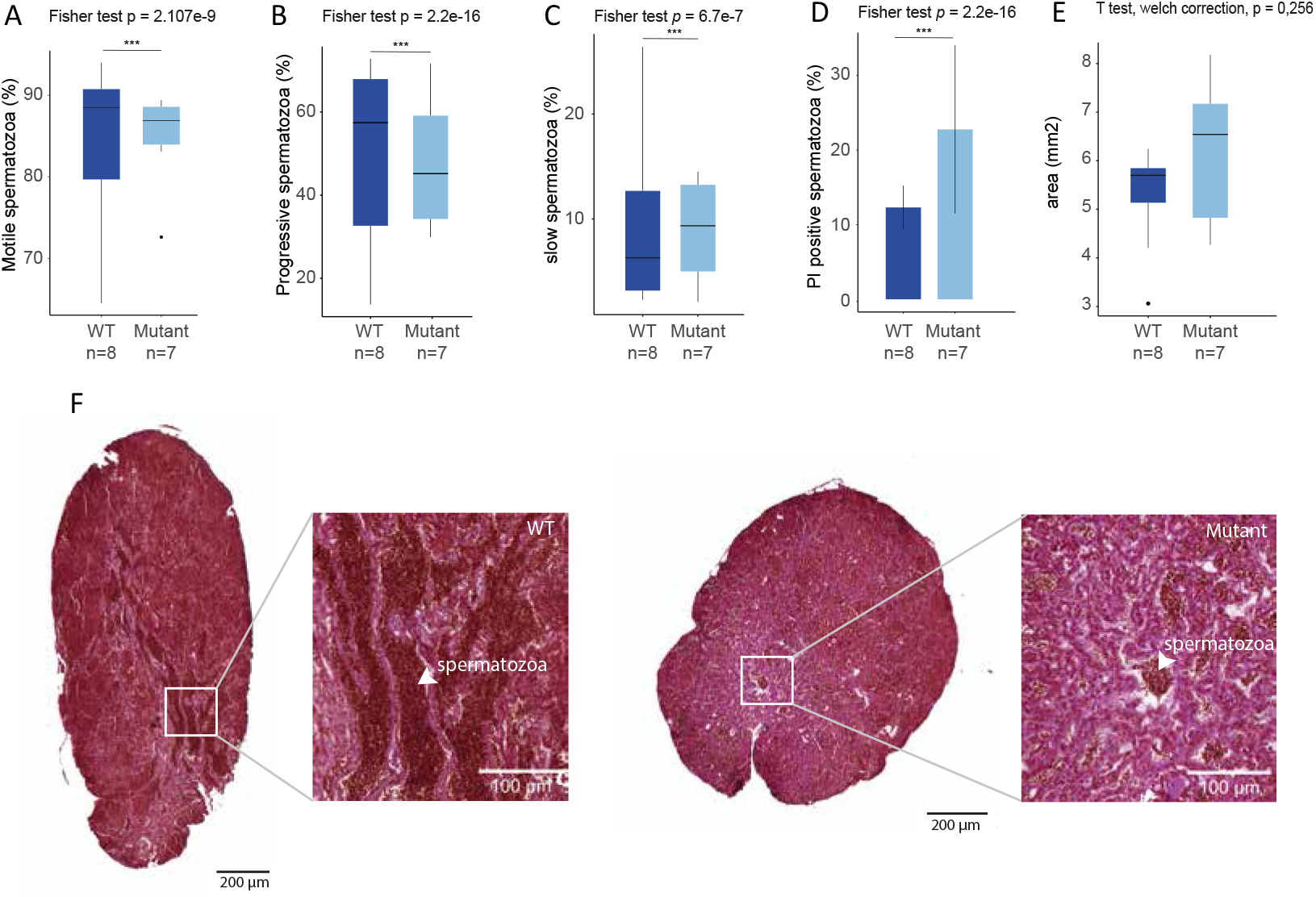
Effect of miR-202-5p target site disruption on testicular development and sperm motility and quality. Percentage of motile (A), forward-moving (B), slow (C), and dead (propidium iodine positive) (D) spermatozoa was assessed in WT and *vgll4b* UTR mutants. Average testicular area on sections (E). Representative pictures of histological sections in WT and mutants (F).

### miR-202-5p target sites are found in the UTRs of both VGLL and TEAD family members cross vertebrate species

In a previous study, we functionally demonstrated that *tead3b* was a phenotypic target (*i*.*e*. target whose de-repression contributes to the phenotypic outcomes) of miR-202-5p involved in the regulation of female fertility in medaka (17). Here we show that *tead3b* is not functionally targeted by miR-202-5p in male medaka. Instead, we demonstrate that *vgll4b*, an antagonist of *tead3b*, is a phenotypic target involved in the regulation of male, but not female, fertility. In teleost fishes, the miR-202-5p binding site in *tead3b* is evolutionarily conserved across species. Even though we initially thought it might have been lost in some species, including zebrafish (17), we were able to identify a miR-202-5p target site in *tead3b* 3’UTR following recent updates of genomic databases (Figure 5). A target site is also present in spotted gar, suggesting its presence in the teleost ancestor. To evaluate the possible long-term conservation of *tead/vgll* targeting by miR-202-5p across species, we analyzed predicted target sites in different fish species, including medaka, zebrafish and spotted gar, and in mice, rat and humans. We identified miR-202-5p target sites in the 3’ UTR of *vgll4b* in medaka, zebrafish and spotted gar, consistent with a conservation of this target site across fishes, similar to *tead3b* (Figure 5). In humans, rat and mice we could not identify miR-202-5p target sites in *TEAD3* or *VGLL4*. However, YAP-dependent gene expression can be mediated by different members of the TEAD family. Similarly, different VGLL proteins can antagonize TEAD-mediated YAP-dependent expression, including VGLL3 (49, 50). In mice, rat and humans, we could identify predicted target sites in both *VGLL3* and *TEAD1* (Figure 5). While the targeting of TEAD1 and VGLL3 by miR-202-5p remains speculative without functional evidence, our results, including the presence of conserved sites across species (Figure 5), shows that these mRNAs are possible targets. Together, our results suggest long-term conservation of antagonistic *TEAD* and *VGLL* targeting by miR-202-5p across vertebrate species that would be consistent with the strong-gonad predominant miR-202 expression (27) and importance of Hippo/Yap regulations (47, 51) in mammalian reproduction.

**Figure 5.**
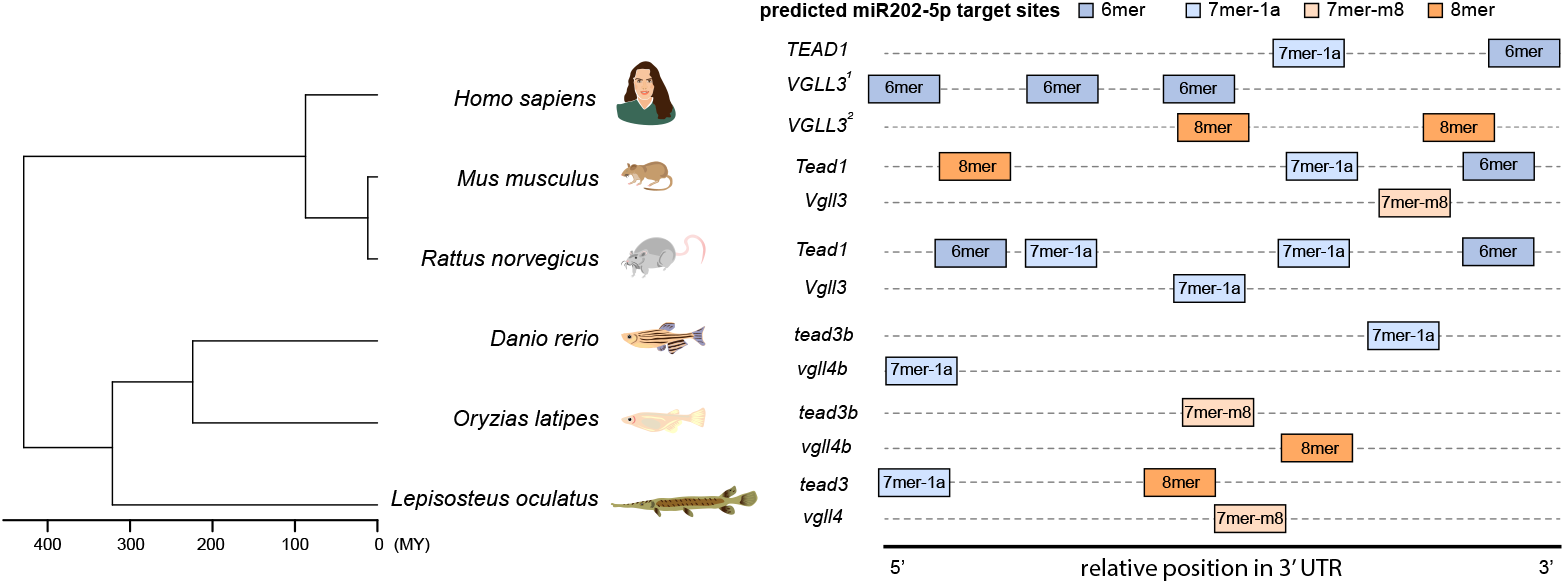
miR-202-5p target conservation in *TEAD* and *VGLL* 3’UTRs across vertebrate species. The type and order target site in the 3’UTR of mRNAs are shown for *TEAD1* (Human: NM_021961,6, Mice: NM_001166584.2, Rat: XM_063267458.1), *VGLL3* (Human: NM_001320493.21^1^ & NM_001320494.2^2^, Mice: NM_001368760.1, Rat: NM_001427793.1), *tead3b* (Medaka: XM_011474938.3, Zebrafish: XM_005166141.5, Spotted gar: XM_006628274.3), and *vgll4b* (Medaka: XM_011474838.3, Zebrafish: XM_073915689.1, Spotted gar: XM_006631022.3).

## Discussion

We identified and functionally validated *vgll4b* as a phenotypic target of miR-202-5p mediating its effect on male fertility in medaka. This was unexpected as miR-202-5p was previously shown to regulate female fertility by targeting *tead3b* and not *vgll4b* (17). To our knowledge, this is the first functional demonstration that a specific miRNA can target two different mRNAs in a sex-dependent manner with major consequences on fertility and, overall, reproductive success. Our results further support the hypothesis that striking phenotypes regulated by a single miRNA gene (1, 2) can be explained by the dysregulation of a single target mRNA (1, 3). In addition, they experimentally support, based on *in vivo* data, the assumption that the biological effects of miRNAs are variable depending on the cellular context, including the quantity and accessibility of theoretical targets. Our results also strongly support the idea that while computationally and *in vitro* predicted targets can give insights into the potential targeting repertoire of a miRNA, they do not provide information regarding the functional relevance of possible targets (4).

It was also unexpected that miR-202-5p targets not only different, but also antagonistic components of the same mechanisms (Yap-dependent transcriptional activity). This is remarkable and these observations demonstrate that opposite gene expression programs are involved in the molecular processes driving male and female gamete formation. Our results show that Yap transcriptional activity plays a pivotal role in gamete formation and that this activity is regulated by miR-202 through the posttranscriptional regulation of Yap partner Tead3b in females and Yap inhibitor Vgll4b in males. Our results are fully consistent with the striking phenotype (low fertility in both males and females) observed when miR-202 is deleted (17, 23) and showcase how a single miRNA can regulate major organism-level phenotypes by leveraging antagonistic transcriptional regulators. While this mechanism was demonstrated in medaka, the strong gonad-predominant expression of miR-202 (26, 29) and the conservation of miR-202-5p target sites of both *tead3b* and *vgll4b* in teleostean and holostean species, including zebrafish and spotted gar, suggest a long-term conservation of this regulatory mechanism in bony fishes. However, the conservation of target sites in TEAD/VGLL in mammals was unexpected, and previously unsuspected. Together, these observations suggest that miR-202, that is also predominantly expressed in both male and female gonads in mammals (27, 28), would also play a major role in gamete formation by targeting *VGLL3* and *TEAD1* in males and females respectively. While this would require functional validation in mammalian models, this hypothesis would be consistent with the numerous experimental evidence providing a functional link between YAP signaling and ovarian function in mammals, including in a pathological context in humans (51).

Here we show that disruption of *tead3b* repression by miR-202-5p leads to a phenotype that resembles polycystic ovarian syndrome (PCOS) at both cellular and molecular levels. This was also unexpected and the dysregulation of key marker PCOS-associated genes demonstrates the similarity of molecular mechanisms involved. Together, the conservation of miR-202 gonad expression and VGLL/TEAD targeting across vertebrates, as well as the striking similarity in key ovarian mechanisms regulating gamete formation suggest a major role of the miR-202/VGLL/TEAD/YAP axis in the regulation of gamete formation and fertility in both males and females across vertebrates. We hypothesize that this axis represents a fundamental regulatory mechanism of both male and female fertility in vertebrates.

## Materials and Methods

### Mutant lines

The mutant lines used in the present study have been established and described in a previous study (17). These lines were designed to obtain either the deletion of the *mir-202* gene (miR-202 KO) or the disruption of the miR-202-5p binding site in the 3’UTR region of tead3b (t*ead3b* UTR) or vgll4b (*vgll4b* UTR).

Genotyping was performed as previously described (17) using the same primers. Briefly, 3-month-old adult fish were anesthetized using a solution of tricaine (225 mg/L; PharmaQ) buffered with sodium bicarbonate (450 mg/mL; Sigma, S5761). A small fragment of the caudal fin was collected for genomic DNA extraction. Fin clips were lysed in 35 μL of alkaline lysis buffer (1.25 M NaOH, 10 mM EDTA, pH 12) by incubation at 95 °C for 45 minutes. The reaction was then neutralized with 35 μL of 2 M Tris-HCl (pH 5). Genotypes were assessed by PCR using primers flanking the targeted deletion sites. Amplification was carried out using JumpStart Taq DNA Polymerase (Sigma-Aldrich, D9307) under the following cycling conditions: initial denaturation at 94 °C for 2 minutes; between 35 to 40 cycles of 94 °C for 30 seconds, 58 °C or 60 °C (depending on the primer pair) for 30 seconds, and 72 °C for 30 seconds; followed by a final elongation step at 72 °C for 2 minutes.

### Reproductive phenotyping

Female reproductive phenotyping was performed as previously described (17). Eggs were collected 2-3 times a week over a 10-week period. Egg collection was performed in the morning, shortly after the onset of the light phase. Spawned eggs were retrieved individually from each female and examined under a stereomicroscope. For each clutch, the total number of eggs was recorded, and embryos were categorized as either developing or non-developing based on their morphological appearance (52).

To assess male reproductive phenotypes, WT and mutant males were mated with WT females. Eggs were collected 2-3 times a week over a 9-week period. Spawned eggs were retrieved individually from each female and examined under a stereomicroscope. For each clutch, the total number of eggs was recorded, and embryos were categorized as either developing or non-developing based on their morphological appearance (52).

### Sperm collection and sperm quality assessment

To reduce sexual behavior and promote the accumulation of spermatozoa in the testes, WT (n = 8) and mutant (n = 7) males of the vgll4b UTR line were isolated for 15 days before collection. Each male was held in a 1.4 L tank with a WT male. To enable individual identification, WT males were subcutaneously marked using fluorescent elastomer tags (Northwest Marine Technology Inc., Shaw Island, WA, USA) (53). For tagging, fish were anesthetized and positioned laterally on a moist surface. The elastomer implant was prepared by mixing the colored polymer and the curing agent in a 10:1 ratio, according to the manufacturer’s instructions. Approximately 0.03 mL of the elastomer mixture was injected subcutaneously beneath the dorsal fin, forming a ∼4 mm-long mark. Following the procedure, fish were placed in a recovery tank before returning to their home tank. Tagged WT males were monitored for one week to confirm retention of the elastomer. Successfully marked individuals were then paired with the males of interest, enabling unambiguous identification of tagged WT versus mutant males during subsequent sperm collection and analysis. After the 15 days isolation period, sperm collection was performed as previously described (54). Male medaka were anesthetized by immersion in a TRIS-buffered MS-222 solution (tricaine methane sulfonate, 225 mg/L) supplemented with sodium bicarbonate (450 mg/L). Once fully anesthetized, fish were gently dried with a paper towel and positioned belly-up (dorsal recumbency) in a holding sponge under a dissecting microscope. Sperm was obtained by applying light pressure along the abdomen in a rostral-to-caudal direction while simultaneously aspirating the expelled milt into collection tubes.

Total motility was analyzed, as previously described (55) using computer-assisted sperm analysis (CASA) and IVOS II system (IMV technologies) using Leja chamber slides (Leja, Netherlands) with a depth of 20 μm (Leja20, four-chamber, LOT: 481815B1). The following CASA parameters were analyzed: (1) total motility (%), slow spermatozoa (%) and (2) progressive motility (%).

### Histological testis analysis

Testes were dissected and immediately fixed in aqueous Bouin’s solution (BOU-OT, Biognost) overnight at 4 °C. Samples were embedded in paraffin and sections of 5 μm thickness were obtained using a microtome (microm HM355S, GMI, Ramsey, MN 55303) and mounted on glass slides using 0.5% albumin (20771.236, VWR) as an adhesive. The sections were dried 2h at 37 °C. For hematoxylin and eosin staining, slides were processed using a Myreva SS-30 automated stainer (MICROM). The program included sequential steps of deparaffinization, staining, and dehydration. Deparaffinization and rehydration steps included successive baths in xylene 10 min (AA16371K7, Fisher Scientific) and rehydration from Ethanol (EtOH) to distilled water (100% 3 min, 96% 3 min, 70% 5 min, distilled water 1 min). For staining, slides were plunged successively into Gill’s hematoxylin force II (CP823, Diapath), a bluing reagent (7301, ThermoFisher Scientific), and aqueous eosin for 2 min 20 s each, with each staining step separated by a 6-min rinse in distilled water. After staining, slides were dehydrated in two successive baths of ethanol at 100°C during respectively 1 min 10s and 2 min 20s and finally cleared in xylene for 10 min (Fisher Scientific). Mounting was performed using a xylene-based medium (SEA-1604-00A, CellPath), and slides were dried for 2 days in an incubator at 37 °C for complete polymerization.

### Quantitative 3D analysis of ovaries

Ovaries were sampled as previously at two key developmental stages corresponding to the end of the juvenile phase and beginning of the reproductive phase (90 dpf) and the peak reproductive phase (160 dpf) when WT females typically reach a plateau in egg production. Ovaries were fixed in situ overnight at 4 °C with 4% PFA in 1X phosphate-buffered saline (PBS; pH 7.4) under gentle agitation. Ovaries were then dissected, rinsed three times in 1X PBS (30 min, 2h, and 2h) at room temperature (RT), and progressively dehydrated in increasing EtOH concentration (25%, 50%, 75% in PBS and 100%, 30 min each at RT). Dehydrated ovaries were stored in 100% EtOH at – 20 °C until further use. Ovaries were gradually rehydrated from EtOH to phosphate-buffered saline at RT under gentle agitation: 75% EtOH 2h, 50% EtOH 2h, 25% EtOH 2h. Samples were then incubated in PBS at 4°C. For tissue clearing, ovaries were incubated in a 1:1 dilution of CUBIC reagent and PBS for 3 h at RT under agitation, followed by full-strength CUBIC reagent for 3.5 days at 37 °C in a hybridization oven. The CUBIC solution was refreshed after the first 24 h. Samples were then rinsed six times in PBS at RT, ensuring thorough rinsing at each step.

For nuclear staining, ovaries were incubated for 2.5 days at 37 °C in a hybridization oven with an anti-Methyl Green dye (323829, Sigma-Aldrich) diluted 1:500 in PBS supplemented with 0.1% Triton X-100 (PBSTx). Following incubation, samples were rinsed five times in PBS containing 0.1% Tween-20 (PBSTw) at RT under gentle agitation and in the dark.

Samples were subsequently dehydrated through a graded methanol series prepared in PBS containing 2% Tween-20, without intermediate PBS storage steps: 25% methanol for 12 h, 50% methanol for 8 h, 75% methanol for 36 h, 98% methanol for 8 h, and finally 100% methanol for 12 h, all at RT and in the dark. Clearing was completed by incubation in ethyl cinnamate (ECi) for 8 h at RT, then transferred to fresh ECi for long-term storage at RT.

Confocal imaging of ovaries from 90 and 160 dpf females was performed using a Leica TCS SP8 laser scanning confocal microscope equipped with a 16×/0.6 IMM CORR VISIR HC FLUOTAR objective (15506533, Leica). Due to the limited working distance of the objective, each sample was imaged in two steps: first with the ventral side oriented toward the objective to image the upper part of the ovary, then flipped with the dorsal side facing the objective to capture the lower part of the ovary—ensuring complete imaging of the ovarian structure, as previously described (56). Images were acquired at a resolution of 512×512 pixels, with a scanning speed of 400Hz for juveniles and 600 Hz for adults (bidirectional), an optical zoom of 0.75, and a z-step of 3 and 6 μm for juveniles and adults, respectively. Line accumulation was set to 2, and frame average to 2. 3D image analysis for follicle segmentation—including image preprocessing, segmentation using deep learning, postprocessing, and follicle diameter quantification—was carried out according to a previously established pipeline (57).

### RNA-seq analysis

To determine the impact of the dysregulation of tead3b by miR-202-5p, we dissected individual ovarian follicles from whole ovaries originating from WT (n = 7) and *tead3b* UTR mutant (n = 11) fish at both early (EV) and late (LV) vitellogenic stages. To standardize ovarian follicular development, ovaries were taken at the same time during the reproductive cycle following ovulation. Ovarian follicles were isolated and selected based on their diameter (EV: 200-300μm, LV: 400-500μm). Pools of 25 to 30 follicles originating for the same were frozen in liquid nitrogen and stored at -80°C until RNA extraction.

Total RNA was isolated from frozen follicle samples using TRI Reagent (Euromedex, TR118). Follicles were mechanically disrupted in Precellys Evolution tubes (CK28 beads, Bertin Technologies) using a Precellys Evolution homogenizer. RNA extraction was then carried out following the manufacturer’s protocol. RNA quantity was assessed with a NanoDrop ND-1000 spectrophotometer (Nixor Biotech), and RNA integrity was evaluated using an Agilent 2100 Bioanalyzer with the RNA 6000 Nano kit. All samples displayed RIN values above 8.5. Purified RNA was stored at −80 °C until library preparation.

cDNA libraries were generated from a minimum of 2 μg of total RNA per sample. Polyadenylated transcripts were enriched using oligo(dT)-conjugated magnetic beads prior to library construction with the Illumina Stranded mRNA Prep Ligation kit. RNA was then chemically fragmented, and first-strand cDNA synthesis was carried out in the presence of actinomycin D to ensure strand specificity. Second-strand synthesis incorporated dUTP, enabling subsequent degradation of the antisense strand during amplification and thus preserving strandedness.

Library quality control included evaluation of fragment size distribution and DNA yield using a Fragment Analyzer (High and Standard Sensitivity NGS kits), followed by library quantification through qPCR on a LightCycler 480 instrument (Roche) before sequencing. Clustering and sequencing were performed on a NovaSeq6000 (Illumina,) using NovaSeq Reagent Kits (Illumina) and the SBS (Sequence By Synthesis) technique. Samples were processed using two lanes of 100-bp single read SP flow cell. More than 934 million reads were obtained with a number of reads per library ranging from 13 to 49 million. Corresponding raw data were deposited in SRA under accession # PRJNA1314461.

Raw read adapter and quality trimming were performed using Trim Galore (version 0.6.6, Cambridge, UK)56 (parameters: --clip_r1 12, --three_prime_clip_r1 2). Trimmed reads were then mapped against Oryzias latipes (ASM223467v1, Ensembl version 114) coding sequence as transcriptome and genome as decoy sequences, and subsequently quantified using Salmon (version 1.8) through Selective Alignment mode based (58). These steps have been gathered in the form of a RNAseq workflow (10.5281/zenodo.15260130, https://forgemia.inra.fr/lpgp/rnaseq).

Genes (pseudo) counts (>=10 reads per gene in at least 4 samples) for Late/early contrasts according to genotype (WT/*tead3b* UTR) normalization and differential expression (samples paired design model) for all samples have been carried out using DESeq2 with default parameters (59).

The WT Early vs WT Late contrast was used to capture the normal transcriptional dynamics underlying follicle growth in WT and tead3b UTR mutants. Differentially expressed genes (DEGs) were filtered to retain only annotated genes with sufficient expression (basemean > 100 in at least one category: WT early, WT late, mutant early, or mutant late) exhibiting an absolute log2 fold change greater than 1, thus focusing on genes likely to have phenotypic relevance. To identify genes whose regulation deviated from the normal WT dynamics in mutants, we selected the most divergent genes using √ (|log2FCmut-log2FCWT|)). This metric quantifies the divergence of gene expression changes in mutants relative to WT.

### miR-202-5p target prediction across vertebrate species

We performed a comparative genomics analysis to identify predicted miR-202-5p target genes in vertebrates, with a particular focus on transcription factors such as *TEAD* and their cofactors *VGLL*. We included tree mammalian species (humans, mice and rat), two model teleost species (medaka and zebrafish) and the spotted gar, an holostean species (a sister group of teleost used to bridge the gap between humans and teleost fishes (60)). Genome assemblies and gene annotations were obtained from NCBI (*Oryzias latipes*: GCF_002234675.1, *Danio rerio*: GCF_049306965.1, *Lepisosteus oculatus*: GCF_040954835.1, *Homo sapiens*: GCF_000001405.40, *Mus musculus*: GCF_000001635.27, *Rattus norvegicus*: GCF_036323735.1) and 3’UTR coordinates were reconstructed from mRNA and CDS features in the NCBI GFF3 files. For each transcript, the genomic region downstream of the last CDS (or upstream of the first CDS on the minus strand) was defined as the 3’UTR, and corresponding nucleotide sequences were extracted with bedtools getfasta. The longest 3’UTR per gene was retained for miRNA target site prediction. miR-202-5p target sites were predicted using TargetScan, which detects canonical 8mer, 7mer, and 6mer seed-matching sites in 3’UTRs. For each species, TargetScan output was parsed to obtain gene identifiers, site type (8mer-1a, 7mer-m8, 7mer-1a, 6mer) and site coordinates. Orthology relationships and ancestral gene information were retrieved from Genomicus (61) and used to link predicted miR-202-5p target sites to teleost orthologues of genes of interest across species. This framework allowed mapping of miR-202-5p sites onto TEAD/VGLL orthologues and other candidate genes to assess the conservation of 3’UTR targeting across species.

## Ethics statement

All experiments were conducted in accordance with EU Directive 2010/63/EU for the protection of animals used for scientific purposes and were approved by the INRAE-LPGP Animal Care and Use Committee (#M-JB012[#1234]).

## Acknowledgments

The authors thank the INRAE fish facility staff for fish rearing and the MGX facility for RNA sequencing. MGX was supported by the France Génomique National infrastructure, funded as part of “Investissement d’Avenir” program managed by Agence Nationale pour la Recherche (ANR-10-INBS-09).

## Funding

This work was supported by Agence Nationale de la Recherche under grant agreement ANR-21-CE20-0023 (MicroHippo) to JB and HS.

## Notes

### Competing Interest Statement

The authors have declared no competing interest.

